# Metabolic Trade-offs can Reverse the Resource-Diversity Relationship

**DOI:** 10.1101/2023.08.28.555123

**Authors:** Zachary R. Miller, James P. O’Dwyer

## Abstract

For species that partition resources, the classic expectation is that increasing resource diversity allows for increased species diversity. On the other hand, for neutral species, such as those competing equally for a single resource, diversity reflects a balance between the rate of introduction of novelty (for example by immigration or speciation) and the rate of extinction. Recent models of microbial metabolism have identified scenarios where metabolic trade-offs among species partitioning multiple resources can produce emergent neutral-like dynamics. In this hybrid scenario, one might expect that both resource diversity and immigration will act to boost species diversity. We show, however, that the reverse may be true: when metabolic trade-offs hold and population sizes are sufficiently large, increasing resource diversity can act to reduce species diversity, sometimes drastically. This reversal is explained by a generic transition between neutral- and niche-like dynamics, driven by the diversity of resources. The inverted resource-diversity relationship that results may be a signature of consumer-resource systems with strong metabolic trade-offs.

## Introduction

Understanding the drivers of biodiversity is one of the central goals of ecology, informing the conservation, restoration, and potentially the design and manipulation of complex ecological communities. Theory based on the differentiation of species emphasizes the role of distinct species niches, for example deriving from specialization on different resources (Chase and Leibold, 2009; MacArthur, 1970, 1972; Tilman, 1982). These differences can act to stabilize the coexistence of diverse communities (Levine and HilleRisLambers, 2009; Schoener, 1974; Tilman, 1982). Diversity in this view is strictly bounded by the number of limiting factors in the environment, a classic result known as the competitive exclusion principle (Levin, 1970; MacArthur, 1970; Schoener, 1974). Thus, a fundamental prediction of niche-based theories is that increasing niche (e.g. resource) diversity should drive increased species diversity (Dal Bello et al., 2021; MacArthur, 1972; Schoener, 1974; Stein et al., 2014). But historically, it has been a challenge to reconcile the competitive exclusion principle with the remarkable diversity of natural communities. Hutchinson famously articulated this problem as the “paradox of the plankton” (Hutchinson, 1959), although the paradox applies equally to tropical rainforests (Wright, 2002), soil microbial communities (Torsvik et al., 2002), and other hyperdiverse ecosystems where many species coexist on apparently few resources or other limiting factors.

Neutral theories, on the other hand, posit that species differences are unimportant for ecological dynamics, and consequently that communities are governed by the stochastic drift of ecologically equivalent species (Chave, 2004; Hubbell, 2001, 2005). Neutral theories predict that the key determinants of diversity are the rate of introduction of novel species (via speciation or immigration) and the total community size (Chave, 2004; Haegeman and Loreau, 2011; Hubbell, 2001; Woodcock et al., 2007). These models evade Hutchinson’s paradox, because diversity can be arbitrarily high given enough input of novelty in a large enough group of individuals. Beyond their consistency with high species richness, simple neutral models have surprising power to explain biodiversity patterns in a range of ecosystems (Bell, 2000; Hubbell, 2001; Volkov et al., 2003; Woodcock et al., 2007). However, these successes are surprising precisely because the equivalence assumption at the heart of neutral theories seems plainly incompatible with well-documented trait differences between species (Purves and Turnbull, 2010), which result in well-characterized differential resource use and habitat associations (D’Andrea et al., 2020b; Schoener, 1974; Webb and Peart, 2000; Zeng et al., 2022), and apparently non-neutral dynamics (Clark and McLachlan, 2003; Fargione et al., 2003; Levine and HilleRisLambers, 2009; Wootton, 2005).

One possible resolution to these tensions is the emergence of neutral-like dynamics and biodiversity patterns in fundamentally non-neutral communities (Gravel et al., 2006; Haegeman and Loreau, 2011; Holt, 2006). Such communities might exhibit neutral-like dynamics for many species at steady-state, while revealing non-neutral responses to large perturbations and differences in ecologically-relevant traits measured “out of context”. Models incorporating both species trait differences and stochastic factors have been used to show how and when neutral drift can dominate niche-driven dynamics (D’Andrea et al., 2020a; Fisher and Mehta, 2014; Gravel et al., 2006; Haegeman and Loreau, 2011). These models demonstrate that neutral-like outcomes can persist when the strict assumption of ecological equivalence is broken, provided that species differences are limited in specific ways. For example, Fisher and Mehta (2014) showed that competitive communities can appear statistically neutral when the variation in species’ interaction strengths falls below a threshold set by the stochasticity of the environment. D’Andrea et al. (2020a) similarly showed that, among consumers competing for resources, neutral-like dynamics arise when variation in the resource preferences of different consumers is sufficiently small. Studies beginning with the work of Scheffer and van Nes (2006) have also shown how “nearly neutral” groups of species might emerge dynamically, as competitive exclusion effectively prunes communities into one or more clusters of similar species, whose trait differences fall below a threshold for statistical neutrality (D’Andrea et al., 2020b; Holt, 2006). Such clusters can arise and persist through a balance of exclusionary and evolutionary processes (Scheffer and van Nes, 2006).

A related possibility is that fitness trade-offs might produce neutral dynamics even in communities with substantial interspecific trait differences, by equalizing per capita demographic rates at steady state (Hubbell, 2005, 2006; Lin et al., 2009; Ostling, 2012). Posfai et al. (2017) recently studied a simple consumer-resource model where the imposition of metabolic trade-offs produces exactly this behavior (see also Erez et al. (2020)). In their model, consumers face an allocation constraint limiting their ability to grow on different substitutable resources, and as a result the community can drive resource availability to a point where any number of consumers have equal fitness. Posfai et al. showed that a deterministic version of this model violates the competitive exclusion principle, although the resulting equilibrium community is not stabilized against all small perturbations, as usually required for stable coexistence. Instead, the model possesses neutral directions along which species abundances can drift, and in a stochastic version of the model, neutral dynamics emerge. This model has inspired significant interest, particularly in microbial ecology, where there has been a productive effort to relate constraints on cellular metabolism to community dynamics (Ferenci, 2016; Harcombe et al., 2014; Litchman et al., 2015; Muscarella and O’Dwyer, 2020; Pacciani-Mori et al., 2021).

Notably, the model of Posfai et al. is similar to other models of emergent neutrality in that neutral dynamics emerge only when consumers are sufficiently similar in a well-defined sense – their total consumption capacity summed over resources. But this notion of similarity is much less restrictive than in the examples above, allowing neutral coexistence of consumers with potentially little or no overlap in their resource preferences.

Models of emergent neutrality raise important new questions. Chiefly, what controls biodiversity in such ecosystems? Does the diversity of resources affect the diversity of consumers, as predicted by niche theories, or is this relationship severed by the emergence of neutral dynamics? How do resource diversity, immigration rate, and system size act together to shape consumer diversity? Using the model of Posfai et al., we explore these questions in a scenario where neutrality emerges from metabolic trade-offs. We find that the interaction of niche and neutral processes can lead to a new and unexpected resource-diversity relationship, where consumer diversity typically decreases with larger numbers of resources, a behavior not predicted by either theory alone. We show that this phenomenon results because communities transition between niche and neutral dynamics depending on the diversity of resources. We then use theoretical results from stochastic geometry to clarify the conditions under which trade-offs may be expected to generate emergent neutrality.

## Models and Methods

### Metabolic trade-off model

We briefly review the model introduced by Posfai et al. (2017), which is a standard consumer-resource model (MacArthur, 1970; Tikhonov and Monasson, 2017; Tilman, 1982) with an added metabolic trade-off. In this model, *m* consumer species compete for *p* resources which are abiotic, substitutable, and externally-supplied at rates ***s*** = (*s*_1_, …, *s_p_*). Each consumer *σ* utilizes resource *i* at a rate *α_σi_≥* 0. Assuming *α_σi_* is proportional to species *σ*’s investment in utilizing resource *i* — for example, its production of enzymes for importing and metabolizing this resource — Posfai et al. modeled an allocation trade-off by imposing the constraint 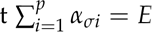, where *E* is a fixed investment budget (e.g. protein synthesis budget) shared by all species. We refer to ***α****_σ_* = (*α_σ_*_1_, …, *α_σp_*) as the metabolic strategy of consumer *σ*, and the set of metabolic strategies allowed under the trade-off as the metabolic simplex. In a sufficiently large system, these dynamics are well-described by the deterministic model

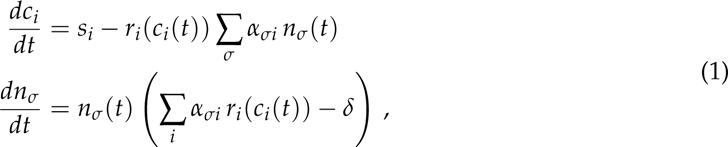

where *c_i_* is the concentration of resource *i* and *n_σ_* is the abundance of consumer *σ*. The rate at which resource *i* is consumed depends on its concentration through the function *r_i_*, which is often modeled as a linear (Type I) or Monod (Type II) function, although the qualitative behavior of the model is unchanged for any *r_i_* that is smoothly increasing and satisfies *r_i_*(0) = 0. All consumers experience mortality (or wash-out from the system) at an equal rate *δ*. We note that an equivalent trade-off would allow consumers to experience different mortality rates and investment budgets, under the assumption that these are proportional (i.e. *δ_i_* ∝ *E_i_*), reflecting the higher metabolic cost of additional investment in consumption (Tikhonov and Monasson, 2017). For simplicity, we focus on the case of equal *E* and *δ* here, and we also neglect resource loss (assuming resource uptake by consumers occurs much faster than resource degradation or outflow) and potential differences in the “investment cost” or “value” of different resources. However, none of these more general (and more realistic) features affect the qualitative behavior of the model, so our main conclusions hold when they are accounted for (see SI Section 1 for additional details).

### Model dynamics

In the absence of the trade-off constraint, no more than *p* consumer growth rates can generally become zero simultaneously, as required for coexistence of the consumers at equilibrium. This is precisely the competitive exclusion principle (Levin, 1970; MacArthur, 1970). The metabolic trade-off, however, implies that when all resources are available in equal concentrations, all consumers have equal per capita growth rates, and, in particular, there is a point where any number of consumers have zero net growth (achieved where *r_i_*= *δ*/*E* for all *i*). Thus, coexistence of *m > p* consumers becomes possible.

In order for coexistence to be realized, though, consumers must also drive the resource rates of change to zero. That is, consumption must balance inflow for each resource. Mathematically, this requires the consumer abundances, ***n*** = (*n*_1_, …, *n_m_*), to satisfy the linear system

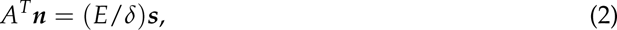

where *A* = (*α_σi_*) is the matrix of consumption rates.

When *m > p*, Eq. 2 is an underdetermined linear system with infinitely many solutions. However, consumer abundances must also be non-negative to be biologically feasible. As noted by Posfai et al., this is the case if and only if the resource supply vector (normalized to (*E*/*S*)***s***, where *S* = ∑*_p_ s_i_*) falls within the convex hull of the *m* consumer metabolic strategies (the points ***α****_σ_*) in the metabolic simplex (see Fig. 1a) ^1^. We refer to this as the *convex hull condition*, and if it is met, then there is a biologically feasible combination of consumer abundances that balances resource inflow. In this case, the dynamics of the ecosystem will approach a point where Eq. 2 is satisfied, *r_i_* = *δ*/*E* for all *i*, and consequently all *m* consumers can coexist indefinitely. If this condition is not met, however, there is no way for equivalent resource concentrations to be sustained — eventually resources will equilibrate at concentrations where *r_i_* differs from *δ*/*E*, and at most *p* consumers will ultimately coexist at equilibrium. Thus, the realized diversity of the community is very sensitive to whether or not the resource supply is contained in the convex hull, particularly when *m* and *p* are very different.

**Figure 1:**
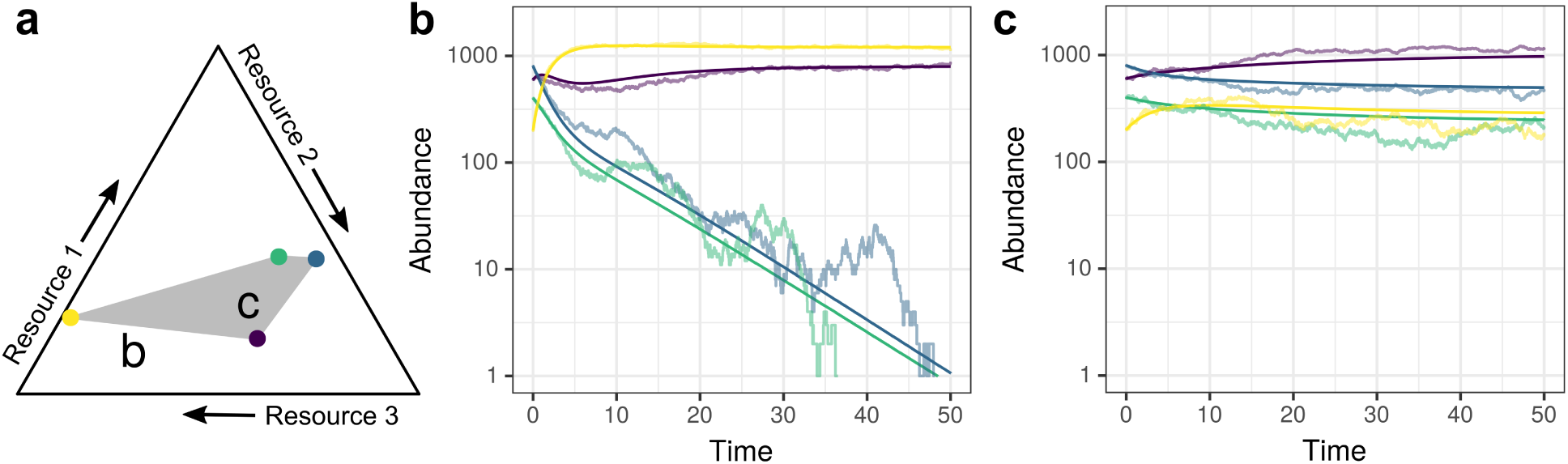
Convex hull condition for the consumer-resource model with trade-offs. Community dynamics depend on whether the resource supply vector falls outside or inside the convex hull of consumer metabolic strategies. (a) Metabolic strategies of four consumer species (colored points) competing for three resources are shown in the two-dimensional metabolic simplex. The convex hull of the consumer strategies is shaded gray. In (b) and (c), we show consumer dynamics corresponding to two different resource supply vectors (indicated by letters in (a)). Deterministic dynamics (solutions of Eq. 1) are indicated by dark lines, and one representative realization of the stochastic BD model is shown with light lines for each scenario. When the resource supply vector is outside of the convex hull (b), two consumer species quickly decline to extinction (at most three — the number of resources — can coexist). When the resource supply is within the convex hull (c), the dynamics approach a neutral manifold where all consumers persist.

Crucially, whenever the convex hull condition above holds, there exists not just one feasible choice of ***n*** that satisfies Eq. 2, but infinitely many, comprising a neutral manifold. The dynamics of Eq. 1 converge to a point on this manifold that depends on the initial conditions. At any such point, the system is stabilized against perturbations that take consumer abundances out of the neutral manifold; the dynamics will quickly return them to it. However, there are *m − p* neutral directions within this manifold that are not stabilized; consumer abundances are free to drift in these directions with no restoring force.

### Stochastic models

Once the neutral manifold is reached, random drift due to stochastic births and deaths of consumers becomes impossible to ignore, even in large systems where the deterministic model provides a very good description of the dynamics away from it. This motivates consideration of a stochastic version of the model. However, we can first simplify the dynamics by observing that, near the neutral manifold, there is a natural separation of resource and consumer timescales, such that resource concentrations typically equilibrate much faster than consumer abundances (SI Section 2; see also D’Andrea et al. (2020a)). Even when the system is far from the neutral manifold, it may be reasonable to assume a separation of time scales between the fast metabolic processes that determine resource concentrations and the slower dynamics of consumer reproduction (Posfai et al., 2017). This allows us to define a stochastic version of the trade-off model (Eq. 1), but tracking only consumer dynamics.

Taking *dc_i_*/*dt* = 0 at all times and letting the quasi-steady-state concentrations *c_i_* be functions of the consumer abundances, the growth-rate function (before mortality) of each species becomes

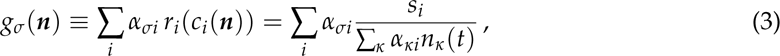

where the denominator is a sum over consumers, giving the total rate at which resource *i* is consumed.

From Eq. 3, the combined growth rate of all consumers is ∑*_σ_ g_σ_*(***n***)*n_σ_* = ∑*_i_ s_i_* = *S*, equal to the total resource supply rate. Therefore we expect the total abundance of all consumers to quickly reach ∑*_σ_ n_σ_* = *S*/*δ*, where growth and mortality are balanced. Taking all of these considerations into account, we define a stochastic birth-death model where the total number of individual consumers is fixed at *T* = *S*/*δ*, which we refer to as the system size. At each time step, a random individual dies and is replaced by the offspring of another individual, chosen through a weighted lottery where species with higher growth rates are more likely to be sampled. More precisely, the probability that the new individual is from species *σ* is given by *g_σ_*(***n***)*n_σ_*/ ∑*_κ_ g_κ_*(***n***)*n_κ_*. Away from the neutral manifold — or in the absence of one — the dynamics of this stochastic process closely track the (consumer) dynamics of Eq. 1, provided *T* is sufficiently large. However, as the system approaches the neutral manifold, where *g_σ_*(***n***) = *δ* for all *σ*, the deterministic dynamics slow to a halt, while the stochastic model continues to drift randomly (Fig. 2). Here, the model exhibits emergent neutrality.

**Figure 2:**
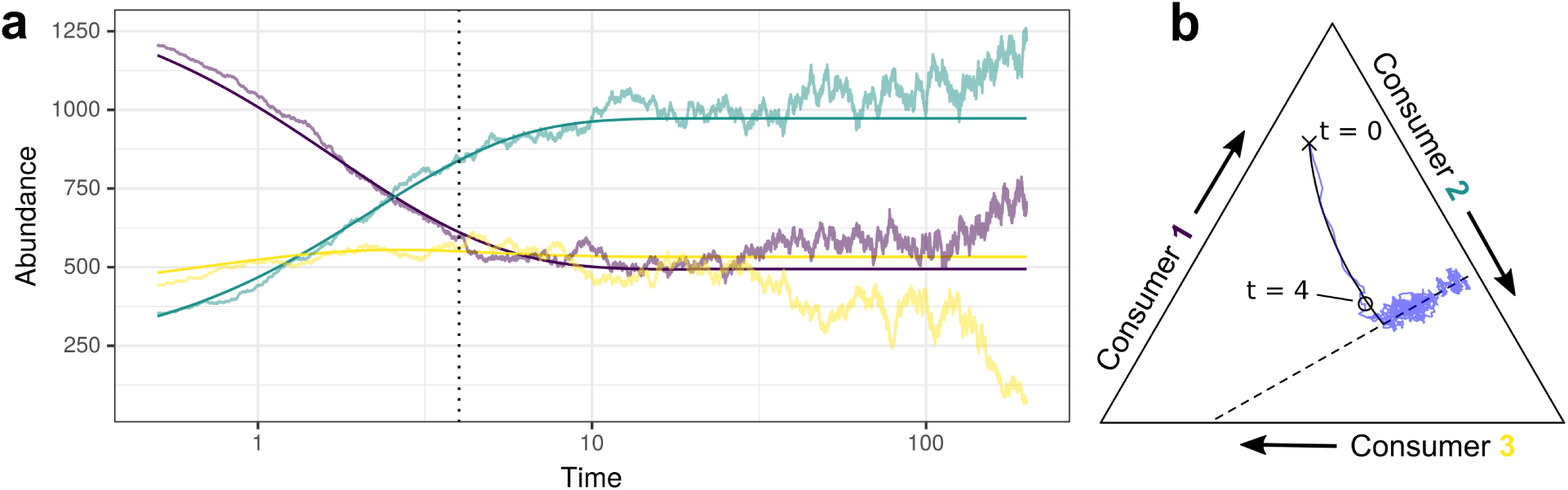
Emergent neutrality in the stochastic (BD) model. When the convex hull condition holds, stochastic dynamics exhibit a transient niche-like phase, followed by neutral drift. (a) An example time series for three consumers competing for two resources illustrates these distinct phases. In the first phase of the dynamics, stochastic trajectories (light lines) closely track the corresponding deterministic dynamics (dark lines) as they approach the neutral manifold; however, once the manifold is reached, the stochastic dynamics drift within its confines, while the deterministic dynamics halt. (b) The same community trajectory visualized in the space of consumer frequencies moves rapidly from the starting point (x) to the neutral manifold (dashed line), at which point the community drifts neutrally along the manifold. To facilitate comparison, the approximate onset of emergent neutrality at time *t* = 4 is indicated in both panels.

This birth-death (BD) model is a discrete, stochastic version of the deterministic model (Eq. 1). The nature of the BD model entails that drift will eventually carry all but one species to extinction. On long time scales, however, diversity can be maintained by the arrival of new species, due to either speciation or immigration from outside the system (henceforth, we refer only to immigration for simplicity). Thus, to investigate steady-state biodiversity we follow Posfai et al. and also consider a version of the model with immigration — a birth-death-immigration (BDI) model, which is exactly like the BD model, except that at each time step the deceased individual may be replaced by a new species, bearing a new metabolic strategy, with probability *ν*. Otherwise, with probability 1 *− ν*, the new individual is sampled from within the community, proportionally to growth rates.

### Numerical simulations

We implemented both stochastic models (BD and BDI) in R (version 4.3.2), following the transition rules outlined above. In simulations where the metabolic trade-off was enforced, all consumer metabolic strategies were sampled uniformly and independently from the metabolic simplex. For simulations with no trade-off, we sampled each consumption rate independently from an exponential distribution with a mean of *E*/*p*. We sampled resource supply vectors in two different scenarios, detailed below. For simplicity and without loss of generality, we fixed *E* = 1 and *δ* = 1 in all simulations and considered different values of *T* (set by *S*) and *ν*. To compare the stochastic and deterministic versions of the model, we also numerically integrated Eq. 1 using a Runge-Kutta method (from the deSolve package in R). To compare our results with a neutral reference model, we simulated the BDI model as described above, but with every individual equally likely to be chosen to reproduce at each time step. To check whether the convex hull condition was satisfied in model communities, we used the geometry package in R.

## Analysis and Results

### Birth-death-immigration simulations

In the BDI model, consumer diversity eventually reaches a dynamical equilibrium where novelty and extinctions are balanced. Thus, we can ask how resource diversity and consumer diversity are related in the BDI model at steady state, with and without metabolic trade-offs. We simulated 50 replicate communities for each case, where each replicate had a fixed resource supply vector (***s***) sampled at random from the simplex and we initialized communities with two consumer species in equal abundances. Each simulation ran for 10^7^ time steps; to investigate steady state communities, we retained abundances from only the last 10^6^ time steps, from which we took 10 community snapshots spaced 10^5^ time steps apart. Replicates and time points were pooled to yield 500 distinct observations for each level of resource diversity. We then summarized consumer diversity in these communities in two complementary ways: by plotting the rank-abundance curve for consumer communities, and by counting the number of consumer species (richness). Results using *T* = 5000 and *ν* = 0.001 are shown in Fig. 3, and other parameter combinations are shown in SI Section 6 (Figs. S2 and S3).

**Figure 3:**
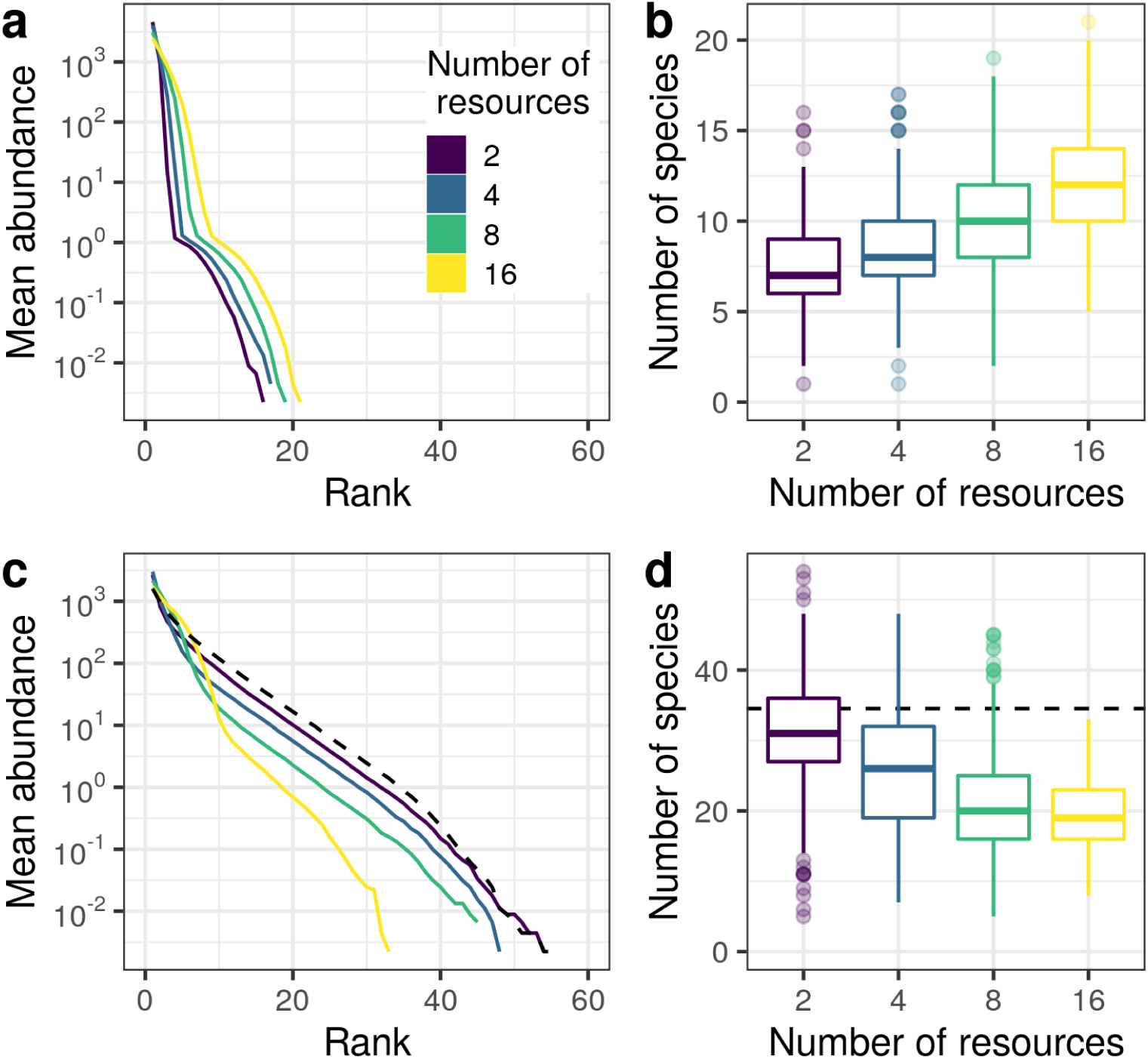
Relationship between resource and species diversity reverses when there are metabolic trade-offs. We plot steady state diversity statistics from the stochastic birth-death-immigration model, simulated over 10^7^ time steps. (a and c) Rank abundance curves for four different levels of resource diversity (averaged over 50 simulations), without (a) and with (c) metabolic trade-offs. (b and d) Distribution of species richness for each level of resource diversity, without (b) and with (d) metabolic trade-offs. Results from a neutral BDI model with the same total community size (*S* = 5000) and immigration/speciation rate (*ν* = 0.001) rate are shown for comparison with the trade-off communities (black dashed lines). When a trade-off constraint is enforced, communities with more resources are typically less even and less rich.

When consumption rates are sampled independently, with no trade-off between them, we find that steady-state biodiversity increases steadily with larger numbers of resources, consistent with classic theory (Levin, 1970; Schoener, 1974). Communities with more resources exhibit flatter rank-abundance curves, indicating a more even composition (Fig. 3a), and higher overall species richness (Fig. 3b).

When we repeat the same simulations with a metabolic trade-off, this pattern is reversed. In this case, communities with more resources typically exhibit steeper rank-abundance curves (Fig. 3c) and fewer total species (Fig. 3d). For the parameters shown in Fig. 3, the median richness with *p* = 2 is more than 50% larger than with *p* = 16 (31 versus 19 species, respectively). For other parameter combinations, this difference can be even greater (Fig. S3f).

Neither niche nor neutral theories can account for this striking inversion of the resource-diversity relationship. Instead, this outcome emerges from the interplay of niche and neutral dynamics in the metabolic trade-off model. Comparing our BDI simulation results against a purely neutral model with the same system size (*T*) and immigration rate (*ν*), we find that the diversity summaries in communities with only 2 resources are almost indistinguishable from the neutral model, while communities with 16 resources appear markedly non-neutral (Fig. 3c-d). This suggests that the dynamics may be more niche- or neutral-like as a function of resource diversity.

### Birth-death simulations

To investigate this idea, we next examined the persistence of diversity in the model without immigration (BD model). In the BDI model, steady state diversity reflects a balance between species lost to extinction and gained through immigration. Because the immigration process has no dependence on the number of resources, we hypothesized that the negative relationship between resource diversity and consumer diversity must arise due to differences in the persistence of species. To isolate the effect of resource diversity on the persistence of consumer diversity, we simulated the BD model with different numbers of resources and a constant number of consumers (*m* = 10) and recorded the time to the first consumer extinction across 3000 replicate communities. As in Fig. 3, we used a system size of *T* = 5000 consumer individuals. The time to first extinction provides a direct measure of how quickly diversity erodes through the dynamics. In these simulations, we vary resource diversity from *p* = 2 to 9, but initial consumer diversity, initial consumer population sizes, and overall resource supply are held constant. We considered two different resource supply scenarios: one where the inflow rates of different resources are chosen uniformly at random from a simplex, with the total resource supply *S* fixed (mirroring our BDI simulations), and a simplified scenario where all resources are supplied at equal rates.

Fig. 4 shows the distribution of times to first extinction for each value of *p* in each of these two scenarios. The results are presented in survival plots, showing the fraction of communities where no extinction has happened up to a given time (the complement of the cumulative distribution function). While there are quantitative differences between the two resource supply scenarios, the qualitative pattern is the same: the time until an extinction is typically longer in communities with fewer resources. This effect is especially pronounced in the random resource supply scenario where, after 50 time units, almost every community with 8 or 9 resources has experienced an extinction, while nearly half of communities with 2 resources remain intact.

**Figure 4:**
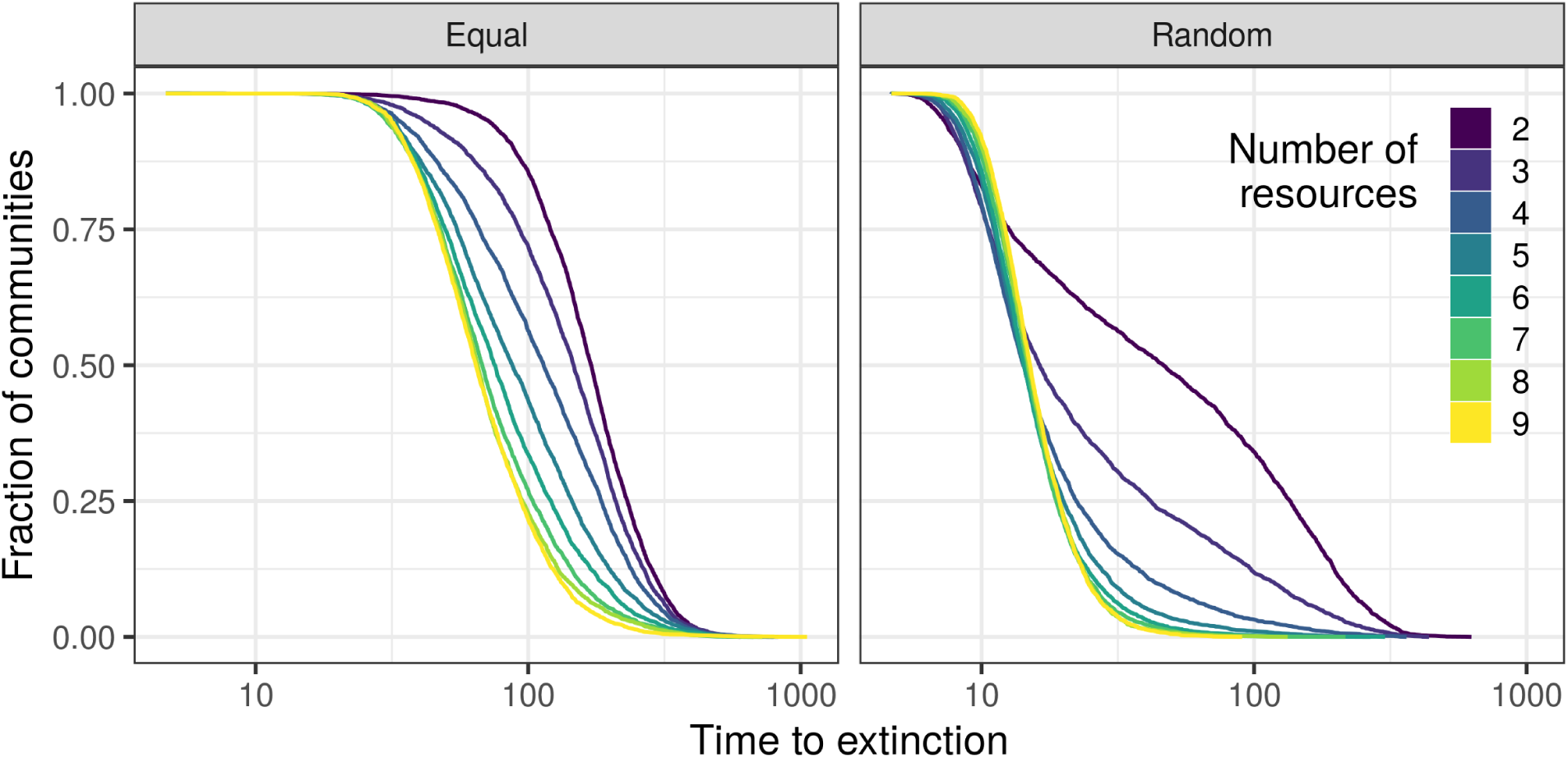
Survival curves show the distribution of times to first extinction in the BD model, for communities with *m* = 10 species and varying numbers of resources (colors). We consider two scenarios for the resource supply: equal supply of all resources (left), or supply chosen uniformly at random from the metabolic simplex (right). Each curve shows the fraction of communities with no extinctions up to the indicated time, summarizing 3000 stochastic realizations for each scenario and number of resources. Times are shown on a log scale. All consumers had equal initial abundances. In both scenarios, communities with fewer resources tend to persist longer before the first extinction.

We expect that the time to first extinction depends strongly on whether dynamics are niche-like or neutral. In the absence of emergent neutrality, a subset of species with lower fitness — corresponding to those excluded in the deterministic version of the model dynamics (Eq. 1) — experience a biased random walk toward zero abundance (as in Fig. 1b). These species will typically be excluded much faster than those experiencing a neutral (unbiased) random walk (Posfai et al., 2017). In the BD model, each community can be classified by whether or not the resource supply vector ***s*** is contained in the convex hull of consumer metabolic strategies, in which case we expect emergent neutrality. In Fig. 5 we plot the conditional times to first extinction, classified in this way. Persistence times are again longer in the equal supply scenario, due to the fact that more equal resource supply promotes more even species abundances. In both supply scenarios, however, it is apparent that extinctions typically occur sooner when the convex hull condition is not met. In other words, some species are lost from the community much more rapidly when there is no emergent neutrality. When the convex hull condition is met, on the other hand, the times to first extinction become much closer to a neutral reference model.

**Figure 5:**
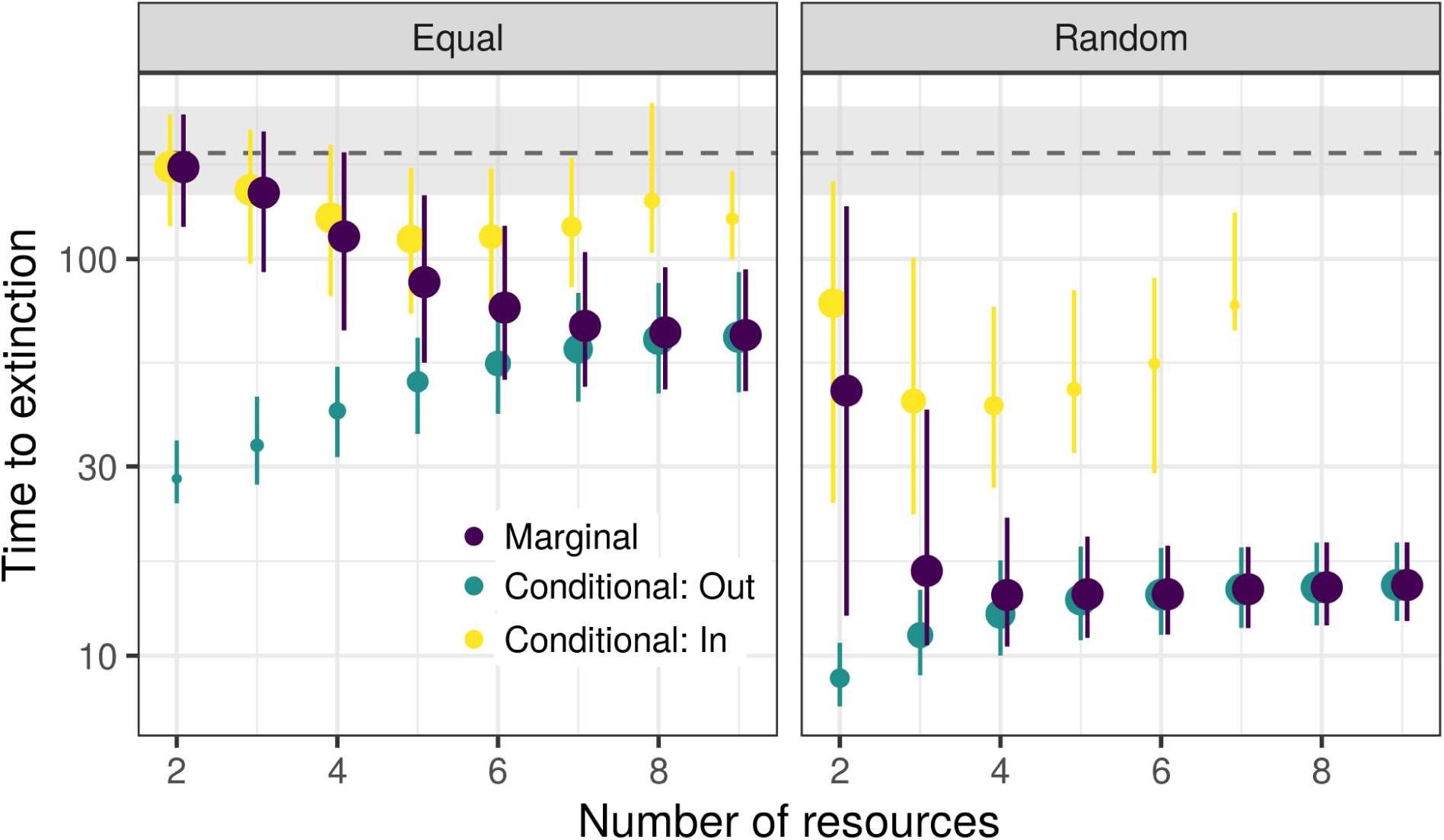
Times to first extinction in the BD model, classified by whether the resource supply is within (yellow) or outside (teal) the convex hull of consumer metabolic strategies. Points indicate the median value for each number of resources; error bars show upper and lower quartiles (note log scale). Point sizes are proportional to the number of observations in each category. For reference, the median (dashed line) and interquartile range (gray) are also shown for a neutral model. When the convex hull condition is met, the distribution of extinction times approaches the neutral expectation; otherwise, extinction times increase as a function of resource diversity. The marginal distribution (purple) shows an emergent negative relationship with the number of resources, resulting from a shift in which outcome is more likely as the number of resources grows.

Notably, the overall negative relationship between resource diversity and consumer persistence disappears once we condition on the convex hull condition. The results within each case — resource supply in or out of the convex hull — conform to neutral and niche theory predictions, respectively. When the resource supply is outside the convex hull of consumer metabolic strategies, we expect niche-like dynamics, where a greater number of resources contributes to consumer coexistence. Indeed, in these cases (teal points in Fig. 5), we find that the time to first extinction is longer with more resources. On the other hand, when the resource supply falls within the convex hull, we expect to find emergent neutrality, and no effect of resource diversity on consumer persistence. As expected, in these cases (yellow points in Fig. 5), we observe little difference in extinction times between communities with different numbers of resources, and no clear trend.

How, then, does the overall negative relationship arise? Because the times to first extinction are much longer under emergent neutrality, regardless of the number of resources, the overall relationship can be explained if communities with fewer resources are more likely to satisfy the convex hull condition. This is exactly what we find. Fig. 5 shows that as the number of resources increases, fewer simulated communities have their resource supply within the convex hull. As a result, the marginal or overall times to extinction, shown in purple, closely track the in-hull results for low resource diversity, but switch to following the out-of-hull results for larger values of *p*. This switch generates an emergent negative relationship between resource diversity and consumer community persistence.

### Convex hull probabilities

Motivated by the dependence of the convex hull condition on resource diversity shown in Fig. 5, we calculated the probability of attaining this condition for various combinations of resource and consumer diversity (*p* and *m*). Specifically, we sampled 1000 sets of consumer metabolic strategies and resource supply vectors for each combination of *m* and *p* shown in Fig. 6, and each supply scenario described above. From these simulations, it is clear that the probability of emergent neutrality decreases with resource diversity very generally. Fig. 6 shows that with 10 consumer species in the equal resource supply scenario, nearly all communities with 2 resources will exhibit emergent neutrality, compared to less than 10% of communities with 8 resources. Emergent neutrality is less likely overall in the random resource scenario, but likewise, we find it more than 75% of the time for *p* = 2, and almost never when *p* = 8. The drop-off is similarly steep for other numbers of consumer species. Thus, communities with few or many resources typically operate in very different regimes: with few resources, the convex hull condition is often satisfied, resulting in emergent neutrality and the potential for higher diversity, while with many resources, the convex hull condition is very unlikely to hold, resulting in niche-like dynamics and fast exclusion of less fit species, despite the presence of the metabolic trade-off.

**Figure 6:**
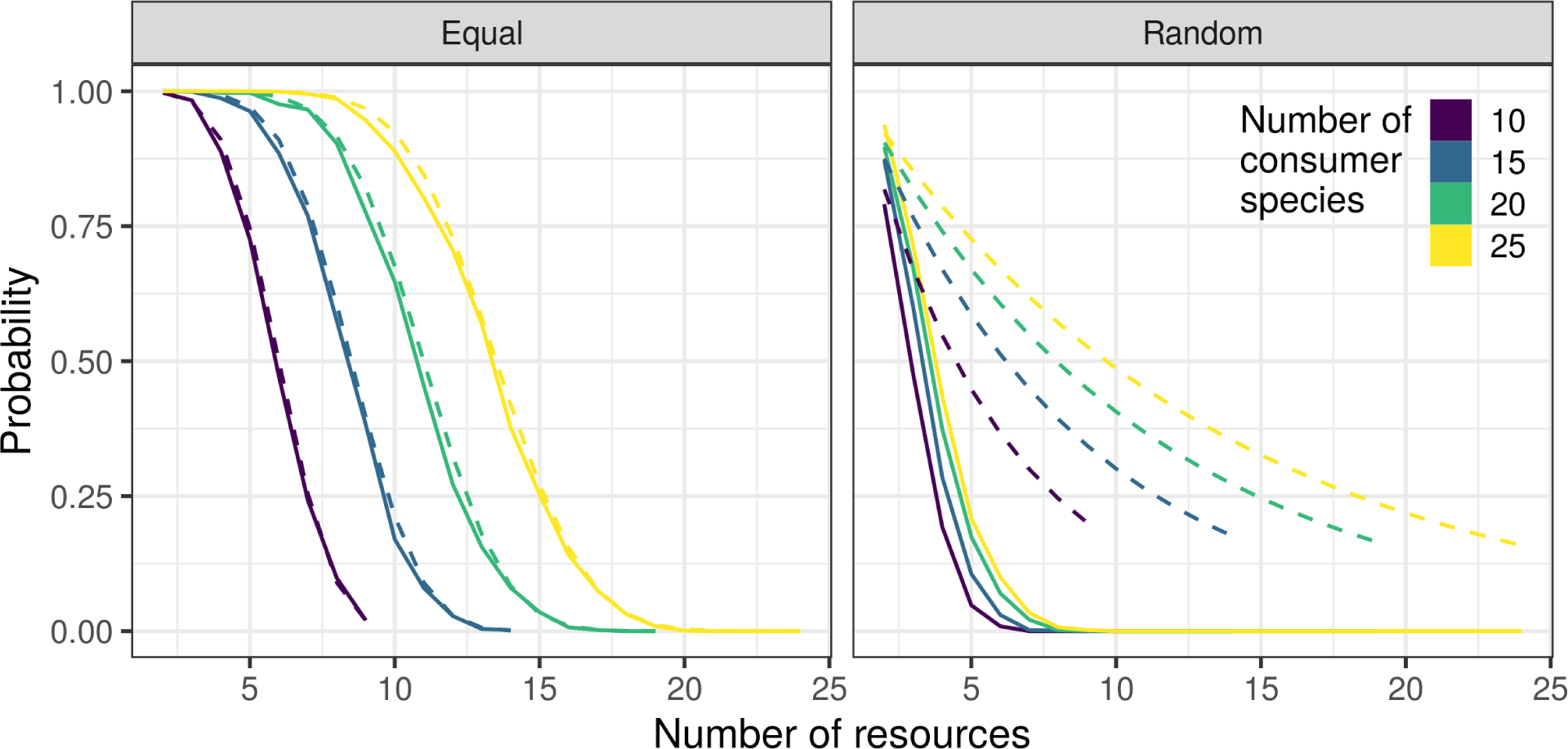
Probability that the resource supply vector falls within the convex hull of consumer metabolic strategies, as a function of the number of resources and consumers. When resources are supplied equally, these probabilities (solid lines) are very well approximated by Wendell’s theorem (Eq. 4; dashed lines). When the resource supply vector is chosen at random, the probability of emergent neutrality is sharply reduced for the same *m* and *p*. These probabilities fall below an exponential upper bound (Eq. 5; dashed lines), which captures the fast initial decline as resource diversity increases. The case *m* = 10 (purple) corresponds to simulations shown in Figs. 4 and 5.

We can take advantage of the geometric formulation of the convex hull condition to better understand how the likelihood of emergent neutrality depends on *m* and *p*. Determining the probability that a given point is contained in the convex hull of some *m* random points in *d*-dimensional space is a well-studied problem in the field of stochastic geometry (in our context, *d* = *p −* 1, the dimension of the metabolic simplex) (Bárány, 2008; Schneider, 2017). A classic result is Wendell’s Theorem (Wendel, 1962), which states that if *m* points are sampled independently from a distribution that is symmetric around a point ***s*** in *p −* 1 dimensions, then their convex hull contains ***s*** with probability

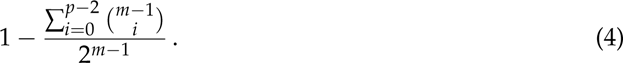

This setting very nearly describes our equal resource supply scenario, where ***s*** lies at the center of the metabolic simplex (Wang et al., 2023). Strictly speaking, the uniform distribution on the metabolic simplex does not satisfy the symmetry assumption in Wendell’s Theorem, but we find that Eq. 4 accurately predicts the probability of finding emergent neutrality in the equal resource supply scenario across range of *m* and *p* values (Fig. 6).

Calculating the same probability for the random resource supply scenario, where ***s*** is also sampled uniformly from the metabolic simplex, turns out to be a substantially harder problem, and only asymptotic approximations or complex integral formulas are known (Affentranger and Wieacker, 1991; Bárány, 2008; Dwyer, 1988; Efron, 1965). However, we can still can derive some insight from Wendell’s Theorem, which is known to provide an upper bound for the probability in this case (in fact, Eq. 4 provides an upper bound for *any* choice of ***s*** and *any* distribution of consumer metabolic strategies, provided they are sampled independently) (Wagner and Welzl, 2001). Additionally, when *p* is substantially smaller than *m*, a better upper bound is given by

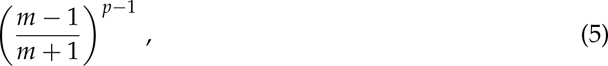

which we derive in SI Section 3 and plot in Fig. 6.

These formulas confirm that, regardless of the resource supply scenario, the convex hull condition (and thus emergent neutrality) is likely to hold when there are few resources compared to the number of consumers (*p ≪ m*), and very unlikely when the number of resources approaches the number of consumers (*p ≈ m*). Furthermore, as *m* and *p* become large, Eq. 4 exhibits a threshold behavior: the probability of finding emergent neutrality in the equal resource scenario drops sharply from close to one for *p < m*/2, to close to zero for *p > m*/2. In the random resource supply scenario, this probability falls off much faster; Eq. 5 shows that it shrinks at least exponentially as *p* increases (Fig. 6). In fact, with many species the random supply scenario also exhibits a probability threshold, but the threshold value is much smaller than *m*/2 (it is proportional to log(*m*)) (Dyer et al., 1992; Frieze et al., 2020). Thus, we should only expect to observe emergent neutrality in this case when there are very few resources, and in general, regardless of how ***s*** and ***α****_σ_* are distributed, Wendell’s Theorem tells us that emergent neutrality is very unlikely when *p > m*/2.

## Discussion

Fitness trade-offs, such as the metabolic trade-offs considered here, may provide a general mechanism by which neutral dynamics can emerge in communities with functionally distinct species. Posfai et al. (2017) demonstrated this possibility using a simple consumer-resource model, motivated by the ubiquitous allocation constraints that govern metabolic strategies in bacteria, phytoplankton, and many other organisms (Ferenci, 2016; Litchman et al., 2015). However, this model can exhibit both niche- and neutral-like dynamics, depending on the specific resource supply and consumer metabolic traits. By investigating a stochastic version of the trade-off model — with and without immigration — we found that niche and neutral dynamics typically arise in different parameter regimes. The interplay of these two regimes can cause a reversal of the positive relationship between resource diversity and consumer diversity predicted by niche theories and the competitive exclusion principle (Levin, 1970; Schoener, 1974; Tilman, 1982).

To understand this outcome, we showed that resource diversity plays a key role in determining the probability of observing niche- or neutral-like dynamics in the model. The fact that this transition is controlled by the number of resources — as opposed to parameters that essentially define niche and neutral regimes, such as the degree of niche overlap or stochasticity — stands in contrast to previous models of emergent neutrality (D’Andrea et al., 2020a; Fisher and Mehta, 2014). Using theoretical results from stochastic geometry, we showed that emergent neutrality is very likely when resource diversity is low, with a sharp transition from typically neutral to typically niche-like dynamics as the number of resources grows relative to the number of consumers. When dynamics are neutral, high diversity can be maintained longer, because no species decline exponentially to extinction. As a result, even though neither dynamical regime exhibits a negative resource-diversity relationship individually, the transition between high-diversity neutral dynamics with few resources and lower-diversity niche dynamics with more resources leads to an emergent negative relationship (Fig. 5).

It is important to recognize that the while the transition from neutral to niche regimes with increasing resource diversity is a very general feature of this model, the resulting relationship with consumer diversity is not necessarily negative. This relationship depends on the typical properties of the two regimes themselves. We find that the relative numbers of resources and consumers sets the dynamical regime (niche or neutral) and the characteristics of the two regimes are controlled by the system size (*T*) and immigration/speciation rate (*ν*) in the neutral case, and also by *p* in the niche case (see Figs. S2 and S3 for simulations across additional parameter values). In the neutral regime, the expected steady-state richness is approximately equal to *−Tν* log(*ν*) (Chave, 2004; Hubbell, 2001). When *T* and/or *ν* are large enough, diversity in the neutral regime can be much higher than in the niche regime, where it is limited by the number of resources, *p*. However, exclusion is not instantaneous in the niche regime, so steady state diversity can sometimes be maintained above *p*, especially if *ν* is large (see, e.g., Fig. S3f). Additionally, in systems with few individuals (*T* small), demographic noise may overwhelm niche-driven dynamics, causing the neutral and niche regimes to appear similar. Both of these factors might act to flatten the resource-diversity relationship. And if *−Tν* log(*ν*) *< p*, diversity may actually be lower in the neutral regime. Overall, we expect that diversity will be substantially higher in the neutral regime, resulting in a negative resource-diversity relationship, when *−Tν* log(*ν*) *> p*, with *T* sufficiently large. More generally, our results provide a foundation for understanding which drivers of biodiversity might be more relevant in communities with metabolic trade-offs, in terms of the system size, immigration rate, and resource diversity.

Empirical studies of resource-diversity relationships have been limited by the challenges of characterizing and manipulating resource diversity, but some relationships from microbial communities have been reported. These include a negative relationship found by Muscarella et al. (2019) between bacterial richness and resource heterogeneity (number of dissolved organic matter components) in lakes. Other experimental studies have reported weakly positive (Dal Bello et al., 2021) or flat (Fu et al., 2020; Pacheco et al., 2021) resource-diversity relationships in microbial communities. These flatter-than-expected relationships might also be consistent with a neutral-niche transition, as explained above. However, it is interesting to note that Muscarella et al. studied a natural gradient of resource diversity, rather than an artificial resource gradient, making it potentially more likely that consumers had evolved to saturate relevant allocation constraints in that environmental context and thus exhibit a trade-off (Bergelson et al., 2021).

Negative relationships have also been reported between environmental heterogeneity and plant diversity (Gazol et al., 2013; Tamme et al., 2010), a context that is closely analogous to the consumer-resource dynamics studied here (Levin, 1970; Miller and Allesina, 2023), and where we expect that trade-offs in performance across different habitat patches could similarly lead to emergent neutrality and neutral-niche transitions. The relationship between plant community diversity and environmental heterogeneity is well studied but variable, with evidence for positive (Lundholm, 2009; Stein et al., 2014), flat (Lundholm, 2009; Tamme et al., 2010), and negative (Tamme et al., 2010) relationships, and substantial evidence that overall resource availability (energy flux) may be a more important driver of diversity than resource heterogeneity (Lundholm, 2009; Stevens and Carson, 2002). Interestingly, our analysis shows that the emergence of neutrality due to trade-offs is one mechanism that might cause consumer diversity to become more sensitive to overall resource supply (*S*, which increases neutral diversity by increasing the number of individuals, *T*) than to the number of resources.

Our theoretical analysis includes several important simplifications. First, we focus on diversity at steady-state, which may require very long times to reach. While we show how differences in steady state diversity arise from differences in extinction dynamics over shorter timescales (Figs. 4 and 5), even these differences might be difficult to observe in short-term experiments. We also assume that consumer metabolic strategies are randomly distributed and resource supply rates are fixed. The former may be unrealistic if consumer strategies have evolved or can be dynamically adjusted to track resource supply (Pacciani-Mori et al., 2020). The latter assumption may be violated if resource availability varies in time, for example when resources are supplied in pulses (Erez et al., 2020; Wang et al., 2023). While a detailed analysis of these scenarios would be more involved, we expect that our broad conclusions are robust to both. Because the convex hull of a set of metabolic strategies “covers” less volume in a higher dimensional metabolic simplex, it should generally be more difficult for consumers to track the supply of many resources at once, and more likely that small changes move the resource supply out of the convex hull. Additionally, Posfai et al. showed that the convex hull condition applies to the time-average of a fluctuating resource supply, implying that our results also characterize this scenario, and that larger convex hulls, which are more likely with fewer resources, increase the probability of finding emergent neutrality in the presence of a fluctuating resource supply, even when consumer metabolic strategies are non-random.

It remains an open question whether the trade-off model introduced by Posfai et al. provides an adequate description of real-world consumer-resource systems. Caetano et al. (2021) have argued that the emergence of neutrality in this model is an artifact of the simple, linear functional form of the trade-off and consumer growth functions. However, in the SI Section 4, we show that emergent neutrality can result from trade-offs in a broad class of consumer-resource models where resources are combined linearly toward growth. Although this condition may sound restrictive, we show that this class of models includes other realistic consumer-resource scenarios, such as multiplicative co-limitation (Harpole et al., 2011; Muscarella and O’Dwyer, 2020; Saito et al., 2008), which are not obviously linear. Another question is whether real-world trade-offs are sufficiently strict to be adequately modeled by an equality constraint (Erez et al., 2020). The qualitative behavior of the trade-off model — including all of our results — is robust to small violations of the equality assumption (Posfai et al., 2017), although sufficiently large variation in species’ investment budgets will result in a loss of the neutral regime (see SI Section 5 for details). Empirical characterization of metabolic trade-offs is an active area of research (Basan, 2018; Ferenci, 2016; Litchman et al., 2015), and closer integration of consumer-resource theory and biophysical constraints could shed light on when and where trade-offs are strict enough to produce emergent neutrality.

We have shown that a transition between neutral and niche dynamics, and an attendant transition between qualitatively different controls on diversity, is a robust consequence of a strict metabolic trade-off. Fundamentally, such a trade-off breaks the typical relationship between resource diversity and consumer diversity because it guarantees a set of resource concentrations at which all consumers have zero net growth. Under these conditions, unless there is some mechanism that couples the number of resources in the system to the number of consumers, such as metabolic crossfeeding (Dal Bello et al., 2021), it is more likely that all resource derivatives vanish simultaneously if there are fewer resources. More broadly, we hypothesize that any tradeoff that equalizes fitness under specific combinations of environmental factors can potentially give rise to emergent neutrality, with a neutral-niche transition as the number of relevant factors grows.

In this study, we focused on the implications of this type of transition for community diversity, finding that it could drive a negative resource-diversity relationship. However, we also note that niche and neutral regimes are characterized by other important differences, such as species turnover through time and stability against perturbations. Further investigation of these other dimensions of neutral-niche transitions may reveal additional signatures and surprising consequences of strong trade-offs.

## Data and Code Availability

All code and simulation data used in this work are available at: https://github.com/zacharyrmiller/tradeoffs reverse resource diversity

## Footnotes

1 The convex hull of a collection of points is the set of all other points that can be obtained as a weighted sum of the original set, where weights are positive and sum to one. In this context, the convex hull defines the set of all resource supply vectors that can be balanced by come combination of consumer strategies weighted by (feasible) consumer abundances.

