## Supplementary Information for "Metabolic Trade-offs can Reverse the Resource-Diversity Relationship"

#### 1 Model simplifications

Following Posfai et al. (2017), we can actually consider a more generic consumer-resource model than the one presented in the Main Text (Eq. 1). Consider the system

$$\begin{aligned} \frac{dc_i}{dt} &= s_i - r_i(c_i(t)) \sum_\sigma \alpha_{\sigma i} \, n_\sigma(t) - \mu_i c_i \\ \frac{dn_\sigma}{dt} &= \rho_\sigma n_\sigma(t) \left( \sum_i v_i \alpha_{\sigma i} \, r_i(c_i(t)) - \delta \right) , \end{aligned} \tag{S1}$$

with $\alpha_{\sigma i}$ subject to the constraints $\sum_{i=1}^{p} w_i \alpha_{\sigma i} = E$. As for Eq. 1 in the Main Text, $c_i$ is the concentration of resource $i$, $n_\sigma$ is the abundance of consumer $\sigma$, $\alpha_{\sigma i}$ is the rate at which consumer $\sigma$ utilizes resource $i$, $\delta$ is the per capita mortality rate of all consumers, and $r_i$ are increasing functions that describe how the utilization rate of resource $i$ depends on concentration. Additionally, we now also incorporate resource loss (degradation or outflow) at rates $\mu_i$, differences in overall consumer growth mediated by $\rho_\sigma$, and varying resource values ($v_i$) and processing costs ($w_i$).

These elaborations on the basic model (Eq. 1 in the Main Text) increase its biological realism and flexibility to describe real-world ecosystems, but we will show that they make little qualitative difference for our analysis.

First, because neutrality emerges when consumer net growth rates are zero, the scaling by $\rho_\sigma$ becomes irrelevant for determining the existence, location, or dynamics around the neutral manifold. However, it is notable that by allowing $\rho_\sigma$ to vary, Eq. 1 can capture trade-offs where $\sum_{i=1}^{p} w_i \alpha_{\sigma i} \neq E$ for all species, but where the mortality rate of each consumer balances its total resource utilization ability (so that $\sum_{i=1}^{p} w_i \alpha_{\sigma i} \propto \delta_i$). Trade-offs of this form have also been investigated, motivated by the assumption that greater investment in resource utilization entails a corresponding metabolic cost (e.g. increased energy demands for respiration) (Tikhonov, 2016; Tikhonov and Monasson, 2017). Our results apply similarly to this implementation of a metabolic trade-off.

Second, Posfai et al. showed that variation in $v_i$ and $w_i$ can be always be "scaled out" of the model, with no loss of generality. In particular, one can define the rescaled metabolic strategies $\boldsymbol{\alpha}'_\sigma = (w_1 \alpha_{\sigma 1}, \dots w_p \alpha_{\sigma p})$ and rescaled resource supply $\boldsymbol{s} = (v_1 s_1, \dots v_p s_p)$. The population dynamics for a system with these rescaled parameters are identical to the dynamics with the original parameters, allowing us to focus on the case where $v_i = 1$ and $w_i = 1$ for all resources. Note, however, that varying these parameters would effectively induce a non-uniform distribution of resource supply vectors in our probabilistic analysis. For additional details on rescaling and simplification of the model, we refer readers to the Supplementary Materials for Posfai et al. (2017).

Finally, considering non-zero resource loss rates ($\mu_i$) requires a minor generalization of the resource equilibrium condition (Eq. 2 in the Main Text). Now we must have

$$A^T \boldsymbol{n} = (E/\delta)(\boldsymbol{s} - \boldsymbol{\mu} \circ \boldsymbol{c}^\star) \tag{S2}$$

where $\circ$ denotes the component-wise product, and $c_i^\star$ satisfies $r_i(c_i^\star) = \delta/E$. Thus, for coexistence, it is no longer the vector $\boldsymbol{s}$ which must fall within the convex hull of consumer metabolic strategies, but the "net resource turnover" vector $\boldsymbol{s} - \boldsymbol{\mu} \circ \boldsymbol{c}^\star$. Notice that if we assume $\mu$ and $r$ are the same for all resources, then $\boldsymbol{\mu} \circ \boldsymbol{c}^\star$ is a constant vector. In this case, our analysis of the equal resource supply scenario is completely unchanged by the consideration of resource loss (assuming $s_i > \mu_i c_i^\star$ for all $i$) while in the random resource supply scenario, resource loss would exert a "de-centering" effect, tending to decrease the probability of attaining the convex hull condition. In the latter scenario, greater resource outflow would pull the distribution of the net turnover vector toward the edges of the metabolic simplex. More generally, resource loss could distort the distribution of net turnover vectors in various ways, but without changing the qualitative features of the model, unless resource loss is large enough that $\mu_i c_i^\star$ exceeds $s_i$, in which case coexistence is no longer possible.

Having seen that the additional complexity of this model changes only quantitative, but not qualitative, details, we focus on a minimal version of the model in the Main Text for clarity. Additionally, we consider a further generalization in SI Section 4, allowing more general forms of metabolic trade-off.

#### 2 Separation of timescales

In the Main Text, we consider a separation of timescales in order to obtain a simplified stochastic model (in particular, a model of consumers only). This timescale separation can be motivated by the fast pace of metabolic reactions and resource uptake, compared to consumer demography, an argument made by Posfai et al. (2017). Here, we show somewhat more rigorously that a separation of timescales naturally emerges around the manifold where emergent neutral dynamics occur. Our analysis, below, makes several simplifying assumptions, but the timescale separation we demonstrate appears to be a more general feature of consumer-resource models near equilibrium (D'Andrea et al., 2020).

Linearizing the model dynamics (Eq. 1 in the Main Text) around a point $(\boldsymbol{c}^\star, \boldsymbol{n}^\star)$ on the neutral manifold yields a matrix differential equation governed by the Jacobian matrix:

$$J = \begin{pmatrix} -D\left(\frac{\partial \boldsymbol{r}}{\partial \boldsymbol{c}}|_{\boldsymbol{c}^\star}\right) D\left(A^T \boldsymbol{n}^\star\right) & -D\Big(\boldsymbol{r}(\boldsymbol{c}^\star)\Big) A^T \\ D\Big(\boldsymbol{n}^\star\Big) A\, D\left(\frac{\partial \boldsymbol{r}}{\partial \boldsymbol{c}}|_{\boldsymbol{c}^\star}\right) & 0 \end{pmatrix}, \tag{S3}$$

where $D(\boldsymbol{z})$ denotes a diagonal matrix with the vector $\boldsymbol{z}$ on the diagonal. On the neutral manifold, we have that $\boldsymbol{r}(\boldsymbol{c}^\star) = \delta/E$ and $A^T\boldsymbol{n}^\star = (E/\delta)\boldsymbol{s}$. Let us also make the simplifying assumptions that $s_i = S/p = s$ for all $i$ (equal resource supply scenario), and that $\frac{\partial r_i}{\partial c_i}|_{c_i^\star} = q$ for all $i$, where $q$ is some positive constant. The latter assumption follows naturally if $r_i = r$ for all $i$; in other words, if all resources are characterized by the same functional response. With these assumptions, the Jacobian becomes much simpler:

$$J = \begin{pmatrix} -\frac{Esq}{\delta}I & -\frac{\delta}{E}A^T \\ qD(\boldsymbol{n}^\star)A & 0 \end{pmatrix}, \tag{S4}$$

where $I$ is the $p \times p$ identity matrix.

The dynamics around the neutral manifold can be decomposed into components corresponding to each eigenvalue/eigenvector pair of $J$. For each eigenpair, the eigenvalue indicates the rate at which a small perturbation decays, while the eigenvalue indicates the direction of the change. The eigenpairs of $J$ are solutions to

$$J\begin{pmatrix} \boldsymbol{u} \\ \boldsymbol{v} \end{pmatrix} = \lambda \begin{pmatrix} \boldsymbol{u} \\ \boldsymbol{v} \end{pmatrix}. \tag{S5}$$

Here, $\boldsymbol{u}$ is a vector with $p$ components, corresponding to the resources, and $\boldsymbol{v}$ is a vector with $m$ components, corresponding to consumers. $\lambda$ is the eigenvalue associated with the eigenvector $(\boldsymbol{u}, \boldsymbol{v})^T$.

First, we notice that if $\boldsymbol{u} = 0$ and $A^T\boldsymbol{v} = 0$, then this provides an eigenvector with a corresponding zero eigenvalue. $A^T$ is $p \times m$ matrix and therefore we have at least $m - p$ linearly independent choices of $\boldsymbol{v}$ such that $A^T\boldsymbol{v} = 0$ (we will assume that the columns of $A^T$ are linearly independent, so that there are exactly $m - p$ linearly independent choices of $v$). Each of these corresponds to a direction within the neutral manifold. As expected, these directions have no components corresponding to resources, because resource concentrations are fixed at equilibrium. However, consumer abundances may be perturbed along these neutral directions with no restoring force ($\lambda = 0$).

To find the other $2p$ eigenpairs of $J$, which must have non-zero eigenvalues, we first use the definition of $J$, above, to write the system

$$-\frac{Esq}{\delta}\boldsymbol{u} - \frac{\delta}{E}A^T\boldsymbol{v} = \lambda\boldsymbol{u}$$
$$qD(\boldsymbol{n}^\star)A\boldsymbol{u} = \lambda\boldsymbol{v}\,. \tag{S6}$$

By substitution we find

$$-\frac{Esq}{\delta}\boldsymbol{u} - \frac{q\delta}{\lambda E}A^TD(\boldsymbol{n}^\star)A\boldsymbol{u} = \lambda\boldsymbol{u} \tag{S7}$$

which can be arranged to yield

$$A^TD(\boldsymbol{n}^\star)A\boldsymbol{u} = \frac{-E}{\delta q}\left(\lambda^2 + \frac{Esq}{\delta}\lambda\right)\boldsymbol{u}\,. \tag{S8}$$

Let $M = A^TD(\boldsymbol{n}^\star)A$ and $\lambda_M$ be an eigenvalue of $M$. The above equation implies that

$$\lambda_M = -\frac{E}{\delta q}\left(\lambda^2 + \frac{Esq}{\delta}\lambda\right)\,, \tag{S9}$$

with $\boldsymbol{u}$ equal to the corresponding eigenvector of $M$ (for each $\boldsymbol{u}$ and $\lambda$, $\boldsymbol{v}$ can be obtained from the relation $qD(\boldsymbol{n}^\star)A\boldsymbol{u} = \lambda\boldsymbol{v}$). Solving this quadratic equation for $\lambda$, we have two solutions corresponding to each eigenvalue of $M$:

$$\lambda = \frac{-Esq}{2\delta}\left(1 \pm \sqrt{1 - 4\frac{\delta^3}{E^3s^2q}\lambda_M}\right)\,. \tag{S10}$$

As $M$ is a $p \times p$ matrix, the $p$ eigenvalues of $M$ clearly generate the $2p$ remaining eigenvalues of $J$.

We have now reduced the problem of characterizing the eigenpairs of $J$ to characterizing the spectrum of $M$. By the definition of $A$ and the assumption that $\boldsymbol{n}^\star$ is feasible, $M$ is a non-negative matrix. Thus, $M$ has a Perron eigenvalue, $\lambda_M^p$, which is largest in magnitude, with a corresponding non-negative eigenvector. Due to the trade-off constraint, which can be written $A\mathbf{1} = E\mathbf{1}$ where $\mathbf{1}$ is a vector or ones, it is easy to see that $\mathbf{1}$ is an eigenvector of $M$ with corresponding eigenvalue $E^2s/\delta$. To verify this, we note

$$A^TD(\boldsymbol{n}^\star)A\mathbf{1} = EA^TD(\boldsymbol{n}^\star)\mathbf{1} = EA^T\boldsymbol{n}^\star = \frac{E^2}{\delta}\boldsymbol{s} = \frac{E^2s}{\delta}\mathbf{1}\,. \tag{S11}$$

This is in fact the Perron eigenpair of $M$, which implies that all $0 \le \lambda_M \le E^2s/\delta$, using the additional fact that $M$ is a positive definite matrix, with all eigenvalues greater than 0.

Now consider the eigenvalues of $J$ corresponding to $\lambda_M^P = E^2 s/\delta$:

$$\lambda = \frac{-Esq}{2\delta}\left(1 \pm \sqrt{1 - 4\frac{\delta^2}{Esq}}\right) . \tag{S12}$$

Typically, we expect the term $\delta^2/(Esq)$ to be small. In particular, we are interested in cases where the total number of consumers, given by $T = S/\delta = sp/\delta$, is large enough so that stochastic fluctuations do not overwhelm the dynamics of interest. For example, in the simulations shown in the Main Text, we have $\delta = 1$, $E = 1$, $s = 5000/p$, and $q = 1$, yielding $\delta^2/(Esq) = p/5000$. Assuming that this term is substantially smaller than 1, $\lambda$ can be approximated using a linearization of the square root:

$$\begin{aligned} \lambda &= \frac{-Esq}{2\delta}\left(1 \pm (1 - 2\frac{\delta^2}{Esq})\right) , \\ \lambda_+ &= \frac{-Esq}{\delta} + \delta , \quad \lambda_- = -\delta \end{aligned} \tag{S13}$$

According to our assumption that $Esq$ is much larger than $\delta$, we have that $|\lambda_+| \gg |\lambda_-|$. In other words, there are two very different timescales associated with $\lambda_M^P$. We can now consider the eigenvector corresponding to each timescale. Recall that $\boldsymbol{u}$ is equal to the Perron eigenvector of $M$, while $\boldsymbol{v} = (q/\lambda)D(\boldsymbol{n}^\star)A\boldsymbol{u}$. For $\lambda_+$, which is the fast rate, $\boldsymbol{v}$ is proportional to $\frac{1}{\lambda_+}$; in other words, the consumer component of this eigenvector is small compared to the resource component. This eigenpair describes perturbations to resources that decay quickly back to equilibrium. For $\lambda_-$, which is the slow rate, $\boldsymbol{v}$ is similarly proportional to $\frac{1}{\lambda_-}$. Now, because $\lambda_-$ is much smaller in magnitude than $\lambda_+$, the consumer component $\boldsymbol{v}$ is large compared to the resource component $\boldsymbol{u}$. In other words, this eigenpair corresponds to slow consumer dynamics near the neutral manifold.

Finally, we note that, because all eigenvalues of $M$ are at least as large as $\lambda_M^P$, the ratio of $\lambda_+$ to $\lambda_-$ is at least as large for all of these eigenvalues. Thus, there are two well-seperated clusters of non-zero eigenvalues of $J$: Those corresponding to $\lambda_+$ for some eigenvalue of $M$, which are large and have corresponding eigenvectors dominated by resources, and those corresponding to $\lambda_-$ for some eigenvalue of $M$, which are small and have eigenvectors dominated by consumers.

#### 3 Derivation of an upper bound for convex hull probabilities

We are interested in bounding the probability that $m$ points (metabolic strategies) in a $p-1$ dimensional simplex contain an additional point (the normalized resource supply vector) within their convex hull, when all $m+1$ points are distributed uniformly and independently. Calculating this probability is straightforwardly equivalent to calculating the expected volume of the convex hull of the $m$ points, or its expected number of vertices. These questions have been studied extensively in stochastic geometry Affentranger and Wieacker (1991); Bárány (2008); Bárány and Buchta (1993); Dwyer (1988); Dyer et al. (1992); Efron (1965); Schneider (2017), but most mathematical results characterize the asymptotic behavior of these quantities as $m$ becomes large. In the regime where $m$ is not much larger than $p$, a better bound can be obtained from elementary considerations.

Notice that in order for some collection of $d$ dimensional points, $\boldsymbol{x}_{1\leq i\leq m}$, to contain a point $\boldsymbol{y}$ in their convex hull, it is necessary that $\min_i(\boldsymbol{x}_i)_j < y_j < \max_i(\boldsymbol{x}_i)_j$ for all $1 \leq j \leq d$. This condition is necessary but not sufficient for $\boldsymbol{y}$ to be contained (consider, for example, the point $\boldsymbol{y} = (1/3, 1/3)$ and the convex hull of $\boldsymbol{x}_1 = (1, 0)$, $\boldsymbol{x}_2 = (0, 1)$, and $\boldsymbol{x}_3 = (1, 1)$ to see that this condition is not sufficient). Using the fact that the points are identically and independently distributed, the probability that $y_j$ is neither the minimum nor the maximum of the $j$th coordinates is $1 - 2/(m+1)$. Then the overall probability that there is no coordinate where $y_j$ is maximal or minimal is simply

$$\left(1 - \frac{2}{m+1}\right)^d = \left(\frac{m-1}{m+1}\right)^d \tag{S14}$$

and setting $d = p - 1$ gives us the upper bound found in the Main Text (Eq. 5).

This bound is exact for $d = 1$ (equivalently, $p = 2$), but becomes looser as $m$ and $p$ grow. In particular, for any fixed $m$, Eq. S14 provides a tighter upper bound than Wendell's Theorem Wendel (1962) for $p$ below some threshold, but for larger $p$ the upper bound due to Wendell's Theorem (from Wagner and Welzl (2001)) becomes better. However, our upper bound clearly highlights the exponential dependence of the convex hull probability on increasing $p$, which leads to a sharp decrease in the probability of emergent neutrality as $p$ grows.

#### 4 More general trade-off models

Posfai et al. (2017) focused on a model of competition for substitutable resources with a simple linear or "sum rule" trade-off. Subsequent analysis has suggested that the resulting dynamics may not be robust to nonlinear trade-offs (Caetano et al., 2021), raising the possibility that these dynamics are biologically unrealistic. Here, we show that emergent neutrality can be produced by a large class of consumer-resource models where resource utilization trades off linearly, and that this class of models includes a surprising diversity of biological scenarios, including, for example, multiplicative co-limitation (Harpole et al., 2011; Muscarella and O'Dwyer, 2020). Furthermore, it is a general feature of these models that emergent neutrality becomes more likely when the number of resources is smaller.

Suppose that the net per capita growth rate of each consumer $\sigma$ takes the form

$$f\left(\sum_i \alpha_{\sigma i} r_i(c_i)\right) \tag{S15}$$

where $c_i$ is the concentration of resource $i$; $r_i$ are increasing functions that determine the "effective" concentration of resource $i$, from the perspective of the consumers; and $f$ is a function that does not depend on $\sigma$. This describes a set of metabolisms where growth for each consumer depends in some arbitrary way on a linear combination of effective resource concentrations. We assume that consumers experience a metabolic constraint of the form $\sum_i w_i \alpha_{\sigma i} = E$. Under these assumptions — and provided that $f(z) = 0$ for some $z$ — there exists a set of resource concentrations, given by $r_i(c_i) = (w_i z)/E$, at which any number of consumers have zero net growth.

Now consider the resource dynamics given by

$$\frac{dc_i}{dt} = h_i(\boldsymbol{c}, \boldsymbol{n}). \tag{S16}$$

The whole system will be in equilibrium if the consumer abundances $\boldsymbol{n}$ satisfy $\boldsymbol{h}(\boldsymbol{c}^\star, \boldsymbol{n}) = 0$, where $c_i^\star$ are resource concentrations that lead to $r_i(c_i^\star) = (w_i z)/E$. As a function of $\boldsymbol{n}$, $\boldsymbol{h}$ is a map from an $m$ (number of consumers) dimensional space to a $p$ (number of resources) dimensional space. When $m > p$, there are in general infinitely many choices of $\boldsymbol{n}$ satisfying the equilibrium condition, provided that there is at least one. Somewhat more precisely, assuming there exists at least one choice $\boldsymbol{n}^\star$ such that $\boldsymbol{h}(\boldsymbol{c}^\star, \boldsymbol{n}^\star) = 0$, and that the Jacobian matrix of $\boldsymbol{h}$ at this point is invertible, then there exists an open set of consumer abundances containing $\boldsymbol{n}^\star$ for which the system is at equilibrium (this is a direct consequence of the implicit function theorem). This set acts as a neutral manifold, along which consumer abundances can drift, exactly as in the model of Posfai et al. Notice that $\boldsymbol{h}(\boldsymbol{c}^\star, \boldsymbol{n}) = 0$ imposes one constraint for each resource on a set of $m$ variables. Thus, with fewer resources there are fewer constraints that must be satisfied, and in general it is more likely that this system can be satisfied — and emergent neutrality attained — when the number of resources is smaller.

As one notable example of a model in this class (along with the model in the Main Text, as well as its close cousin, the MacArthur consumer-resource model with biotic resources), consider the case where

$$\begin{aligned} f(x) &= \exp(x) - \delta, \\ r_i(x) &= \log(x) \quad \text{for all } i, \\ h_i(x, y) &= s_i - \sum_\sigma \alpha_{\sigma i} y_i \prod_\beta x_\beta^{\alpha_{\sigma\beta}} - \mu_i x_i \quad \text{for all } i. \end{aligned} \tag{S17}$$

Then, following Eq. S15, the dynamics of the corresponding consumer-resource system are governed by

$$\begin{aligned} \frac{dc_i}{dt} &= s_i - \sum_\sigma \alpha_{\sigma i} n_i \prod_\beta c_\beta^{\alpha_{\sigma\beta}} - \mu_i c_i \\ \frac{dn_\sigma}{dt} &= n_\sigma \left( \exp\left( \sum_i \alpha_{\sigma i} \log(c_i) \right) - \delta \right) = n_i \left( \prod_i c_i^{\alpha_{\sigma i}} - \delta \right) \end{aligned} \tag{S18}$$

with the constraint $\sum_i \alpha_{\sigma i} = E$ for all $\sigma$. This is a model for multiplicative co-limitation (Harpole et al., 2011; Muscarella and O'Dwyer, 2020; Saito et al., 2008), precisely of the form studied by Butler and O'Dwyer (2020). When all $c_i = \delta^{1/E}$, all consumers have zero net growth and there is a neutral manifold, which is defined exactly as in Eq. S2.

### 5 Relaxing the trade-off constraint

Posfai et al. (2017) considered a "hard" metabolic constraint (an exact equality), which is necessary for the existence of a neutral manifold (Cui et al., 2020). If the total investment budget ($\sum_i \alpha_{\sigma i}$) varies between consumer species, no more than $p$ species can persist at equilibrium. However, if species vary only slightly in their investment budget (a "soft" constraint (Cui et al., 2020; Tikhonov and Monasson, 2017)), then there will be a region of the phase space characterized by very slow consumer rates of change. In effect, the neutral manifold may be replaced by a "nearly-neutral" manifold or "ghost attractor" (Hastings et al., 2018). If rates of change in this region are sufficiently low, then high diversity may persist transiently over long time scales and, in the stochastic birth-death-immigration model, steady state diversity may remain high.

Posfai et al. considered this possibility and derived a rough threshold for how far the trade-off constraint can be relaxed before BDI dynamics become substantially non-neutral. They assumed that resource utilization rates can be modeled by an exact trade-off (i.e. $\alpha$s such that $\sum_i \alpha_{\sigma i} = E$) plus some Gaussian noise, so that a species $\sigma$ has the metabolic strategy

$$(\max(\alpha_{\sigma 1} + \zeta_{\sigma 1}, 0), \ldots, \max(\alpha_{\sigma p} + \zeta_{\sigma p}, 0)) \tag{S19}$$

where $\zeta_{\sigma i}$ are iid Gaussian random variables with mean zero and standard deviation $\Sigma$. By comparing the timescales of the first extinction in this scenario (on the order of $\frac{T\log(T)}{\Sigma}$) versus a scenario with a hard metabolic constraint (on the order of $T^2$), Posfai et al. argued that these timescales — and thus the qualitative dynamics of the two model scenarios — converge when $\Sigma$ reaches a threshold proportional to $\log(T)/T$. For $\Sigma$ below this threshold, the ecosystem will display neutral-like or niche-like dynamics depending on whether or not the convex hull condition is satisfied, exactly as for the model with hard metabolic constraints.

Fig. S1 shows the same birth-death-immigration scenario as Fig. 3 in the Main Text, but where the trade-off constraint is relaxed by adding Gaussian noise with $\Sigma = 0.0005$. Qualitatively, the results are unchanged by this relaxation.

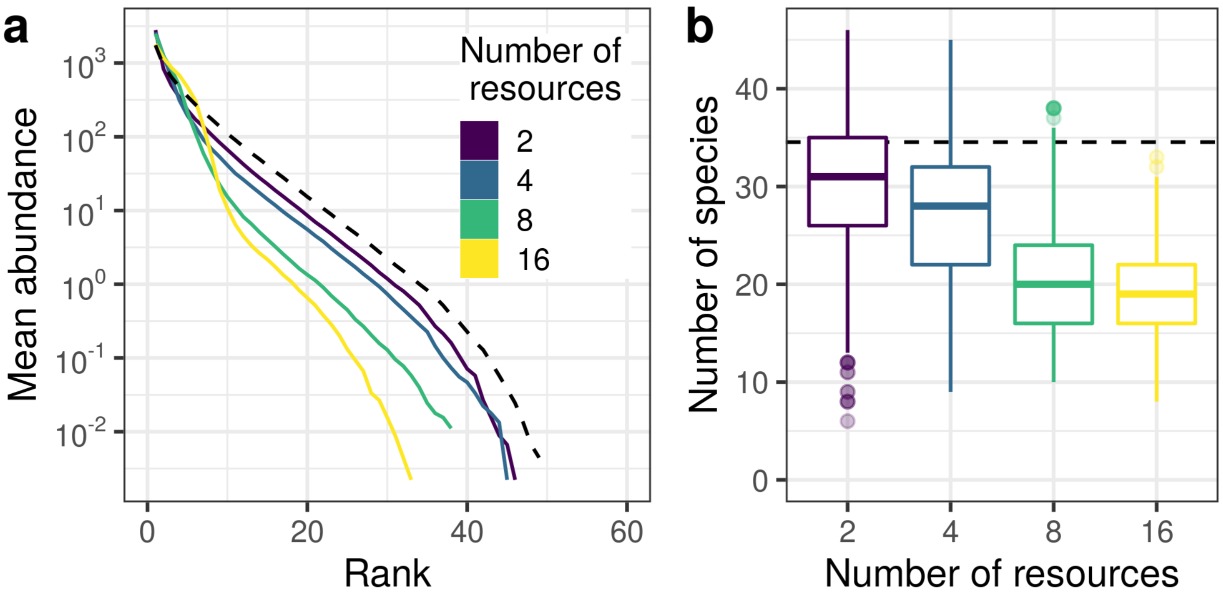


Figure 1: Higher resource diversity leads to lower steady-state consumer diversity in the stochastic birth-death-immigration model with a relaxed trade-off ($\Sigma = 0.0005$). (a) Rank abundance curves for four different levels of resource diversity (averaged over 50 simulations). (b) Distribution of species richness for each level of resource diversity. Results from a neutral BDI model with the same total community size ($T = 5000$) and immigration/speciation rate ($\nu = 0.001$) rate are shown for comparison (black dashed lines). As for the simulations in the Main Text with a hard trade-off, we find that with more resources, communities are less even and less rich.

#### 6 BDI simulations: additional parameterizations

Here, we show results for four different BDI simulation scenarios, as in Fig. 3 in the Main Text but with varying system size, $T$, and immigration rate, $\nu$. In particular we consider values of each parameter that are half and double the values shown in the Main Text, used in all combinations (i.e. $T \in \{2500, 10000\}$ and $\nu \in \{0.0005, 0.002\}$).

Community diversity statistics, summarized by rank-abundance curves and richness, are shown in Figs. S2 and S3, respectively. We observe that, in all cases where the trade-off is imposed, the diversity statistics of communities with two resources closely approximate a corresponding neutral model (i.e., a neutral model with the same $T$ and $\nu$, which control the properties of the neutral communities). This is consistent with our finding that the emergence of neutrality depends primarily on the relative numbers of resources and consumers, while the properties of the emergent neutral regime are determined by $T$ and $\nu$.

In the first two scenarios (a-d in each figure), we used parameter combinations that produce similar neutral richness to the parameters used in the Main Text (neutral model richness is approximately equal to $-T\nu\log(\nu)$). Consequently, we observe that these simulations very closely resemble the results shown in Fig. 3 in the Main Text, despite the variation in parameter values. This suggests that $T$ and $\nu$ primarily act through the composite parameter $-T\nu\log(\nu)$. In the third scenario (e-f in each figure), both $T$ and $\nu$ are larger than in the Main Text, producing even higher diversity in the neutral regime, and a more dramatic negative resource-diversity relationship. Finally, in the last scenario, both $T$ and $\nu$ are smaller than in the Main Text. For these parameter values, neutral richness is low enough ($< 10$ on average) to be comparable to diversity in the niche regime with 16 resources. As a result, we observe a flat resource-diversity relationship in this case.

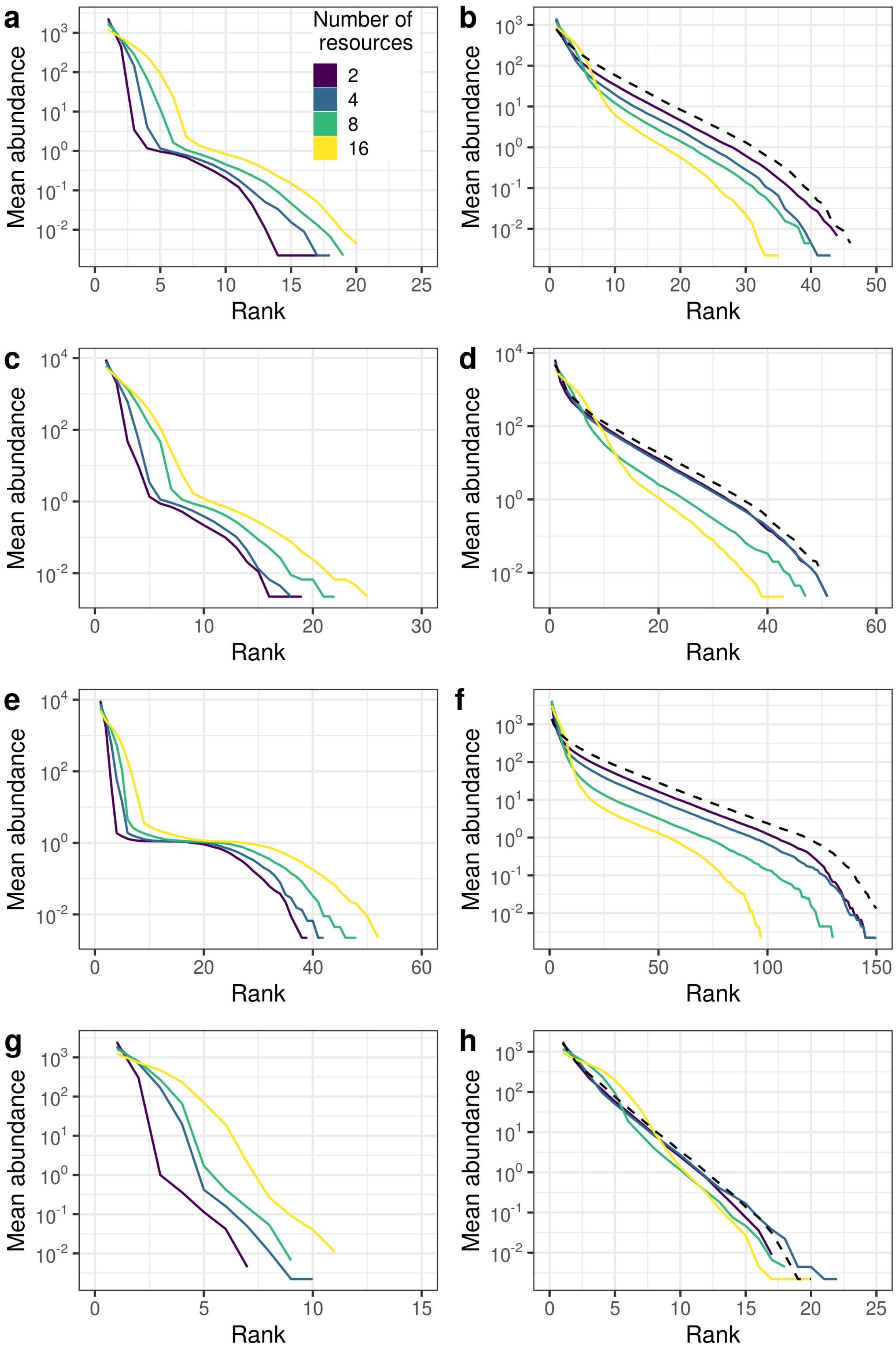




Figure 2: Caption next page.

Figure 2: (Previous page.) Rank-abundance curves for birth-death-immigration simulations. As Fig. 3 in the Main Text, but with varying values of $T$ and $\nu$. For each combination of parameters (rows) we show results without (left) and with (right) a trade-off imposed. Parameter combinations are (a-b) $T = 2500$ and $\nu = 0.002$, (c-d) $T = 10000$ and $\nu = 0.0005$, (e-f) $T = 10000$ and $\nu = 0.002$, (g-h) $T = 2500$ and $\nu = 0.0005$. Results from a neutral reference model are shown with black dashed lines. Note that for the third combination ($T = 10000$ and $\nu = 0.002$), reaching steady state required each simulation to run twice as long ($2 \times 10^7$ time steps). As for all other simulations, we retained the last $10^6$ time steps from each replicate and plot results from 10 equally spaced time points.

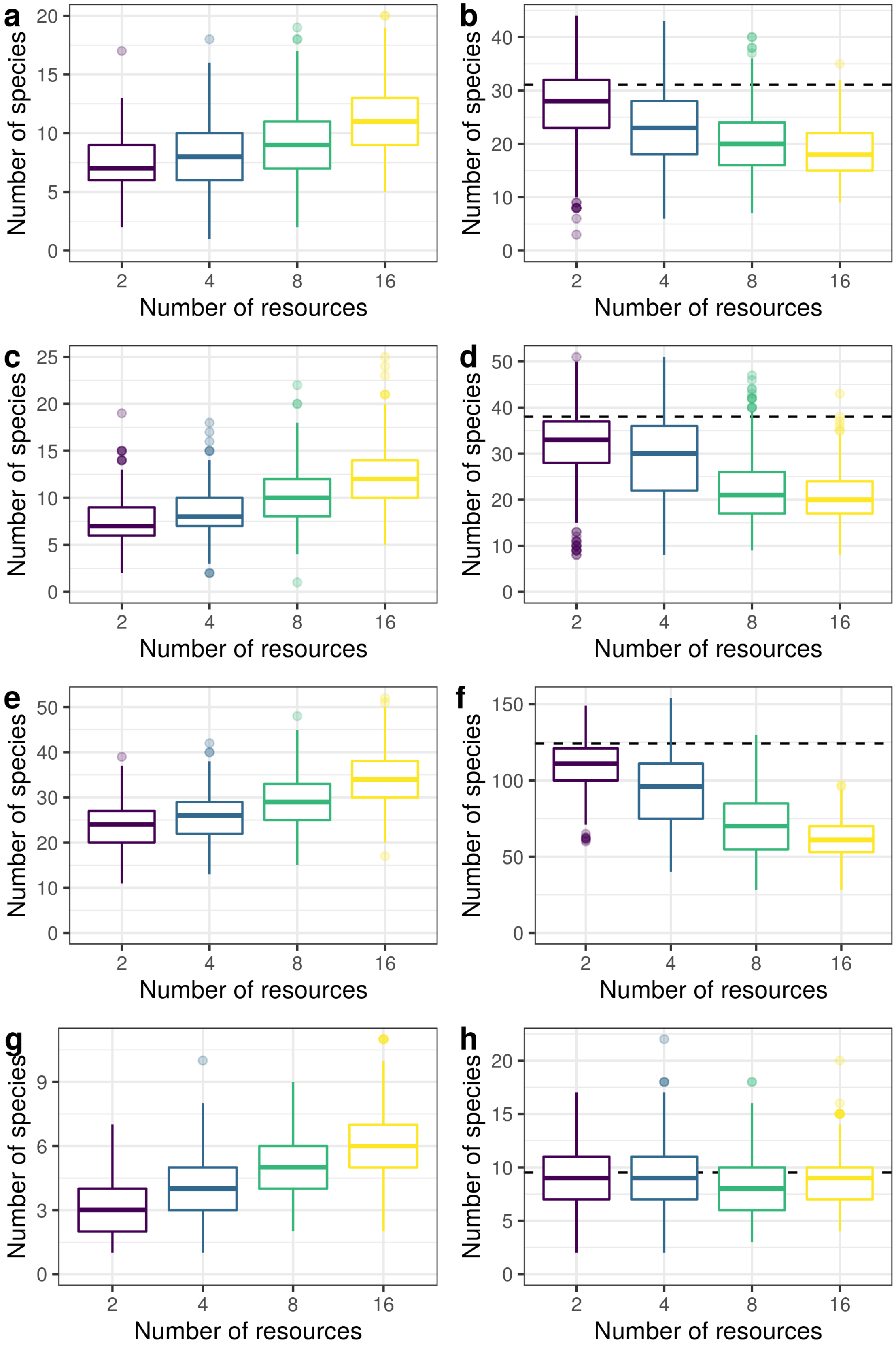




Figure 3: Caption next page.

Figure 3: (Previous page.) Community richness for birth-death-immigration simulations. As Fig. 3 in the Main Text, but with varying values of $T$ and $\nu$. For each combination of parameters (rows) we show results without (left) and with (right) a trade-off imposed. Parameter combinations are (a-b) $T = 2500$ and $\nu = 0.002$, (c-d) $T = 10000$ and $\nu = 0.0005$, (e-f) $T = 10000$ and $\nu = 0.002$, (g-h) $T = 2500$ and $\nu = 0.0005$. Results from a neutral reference model are shown with black dashed lines. Note that for the third combination ($T = 10000$ and $\nu = 0.002$), reaching steady state required each simulation to run twice as long ($2 \times 10^7$ time steps). As for all other simulations, we retained the last $10^6$ time steps from each replicate and plot results from 10 equally spaced time points.
